# Edible fire buffers: mitigation of wildfire with multifunctional landscapes

**DOI:** 10.1101/2021.08.30.458294

**Authors:** Xiao Fu, Abigail Lidar, Michael Kantar, Barath Raghavan

## Abstract

Wildfires ravage lands in seasonally-dry regions, imposing high costs on infrastructure maintenance and human habitation at the wildland-urban interface (WUI). Current fire mitigation approaches present upfront costs with uncertain long-term payoffs. Instead, we show that a simple landscape intervention on human-managed wildlands – buffers of a low-flammability crop species such as banana irrigated using recycled water – can mitigate wildfires, produce food profitably, and provide additional ecosystem services. Recreating a recent, major fire in simulation, we find that a medium-sized banana buffer decreases fireline intensity by 96%, similar to prescribed burns and mechanical thinning combined, and delays the fire by 316 minutes, enabling safe and effective firefighting. We find that under climate change, despite worsened fires, banana buffers will still have a protective effect. We also find that banana buffers with average yield could produce a profit of $56k USD/hectare through fruit sales, in addition to fire mitigation and other benefits.

## Introduction

Sustaining human habitation in seasonally-dry regions of the world is an intensifying challenge with urban expansion and climate change [1]. Anthropogenic temperature increases and consequent global aridity [2] have increased wildfire risk; wildfires nearly doubled in frequency in the Western United States from 1984 to 2015 [3]. Further, climate change is threatening global food production not only for staple grain crops, but also for important fruits and vegetables [4]. Climate change is decreasing food production in existing agricultural regions while increasing fire risk [5], providing an opportunity to simultaneously address both challenges.

Simultaneously, the wildland-urban interface (WUI) – “the area where houses are in or near wildland vegetation” – has increased over 40% in the United States, along with fire risk [6]. WUI land often has high or extreme fire risk [7], with such risk amplified by climate change [8]. Despite this risk, the WUI is increasingly desirable for building structures. Wildfire in the United States imposes annual costs of 70–300 billion USD [9]. California in particular faces major wildfire risk, threatening more than four million homes [10].

There is growing recognition that no one technique will sufficiently mitigate the risk of fire at WUIs. Instead, many techniques must be employed, including: 1) enhanced building and land-use codes, 2) fuel reduction in wildland and WUI areas, 3) increased firefighting resources, and 4) creation and maintenance of firebreaks. Despite this wealth of options, existing strategies have uncertain long-term payoffs with large up-front costs [11, 12].

Choice of vegetation surrounding WUIs is key, but standard “fire resistant” plants do not yield useful outputs [13]. Unlike prior work, here we study the creation of buffers of crop plants on human-managed wildlands that abut the WUI, to mitigate fire spread while producing profitable yields. We focus specifically on non-intermix WUI zones; intermix areas lack a clear boundary for such buffer plantings (here a *buffer* is an area of human-managed wildland that abuts the WUI, with low-flammability vegetation).

### Multifunctional Landscapes for Fire Mitigation

We consider the effectiveness of alternative fire buffers designed for multifunctionality, with careful consideration of the species and their effectiveness for fire mitigation, ease of propagation, ease of maintenance, and yield of high-value outputs to recoup investments. We specifically consider the use of increasingly-available recycled water in these regions.

Planted fire buffers must have high water content at all times [14]. Irrigated orchards and vineyards have insufficient moisture to adequately slow or stop fire spread [15, 16]. Other types of irrigated agriculture are not suited to uneven terrain typical of such high-fire-risk areas. Non-irrigated fire buffers are unlikely under climate change-driven drought conditions to have sufficient water content [17]; the prominent exception is some succulent fire buffers which may be effective but do not produce high-value outputs. Crops such as banana have very high water content, from 93–99% in good moisture conditions to 76–88% under drought [18, 19].

We consider several criteria for potential multifunctional fire buffer crops: 1) minimal management needs, 2) suitability in present and future climates, and 3) low flammability [20]. Such edible fire buffers are thus new agroecosystems in wildland areas that abut the WUI in which the fuel and management is fundamentally different from current circumstances.

## Results

To understand the potential for edible fire buffers to mitigate risk at the WUI, we simulated a historical fire. California has faced numerous major wildland and WUI fires in recent years, yet many historical fires yielded inadequate or inappropriate data for analysis: some progressed too quickly in intermix (e.g., Camp Fire, 2018), others were at a WUI but were suppressed by extensive firefighting (e.g., Getty Fire, 2019), while still others had minimal effect on populated areas. To balance these factors, we selected the 2017 Tubbs Fire. The Tubbs Fire fits the context of the intervention we explore: location in a semi-arid or Mediterranean region, origination in wildland, progression due to prevailing winds through the WUI, and rapid advancement that overwhelms firefighting resources leading to significant loss of life and structures. Additionally, simulations allowed us to illustrate dramatic containment differentials, since the Tubbs Fire caused the most damage at the WUI of any non-intermix WUI fire in California history. Given the fire’s location in Sonoma County and proximity to Napa County, we were also able to compare banana buffers with pre-existing alternatives: vineyards and orchards.

### Replicating the Tubbs Fire

Using FARSITE [21], we replicated the Tubbs Fire in simulation. FARSITE is designed for wildland fire simulation, which is well suited to our context as edible fire buffers involve a change of fuels on human-managed wildlands. No fire simulation tools are well adapted to fires within the WUI and urban areas. In our context, however, the primary analysis is squarely on wildlands, for which FARSITE was designed, as edible fire buffers involve a fuel change of such human-managed wildlands in which a fire may progress toward habitated areas such as in the case of the Tubbs Fire.

As we are also concerned about WUI impacts, not simply total wildfire spread, we separately verified the replication of our simulation on the urban perimeter and the total perimeter relative to ground-truth USGS satellite data [22]. For the burn area in the WUI and urban land in the Tubbs Fire we achieved *F*_1_ scores of {0.68, 0.74, 0.64} with respect to the three available satellite timepoints and {0.75, 0.78, 0.74} for the total area burned, exceeding the scores of general fire models [23], despite the fact that FARSITE [21] is primarily intended for wildland fires. Our focus in this replication is on the area of WUI/urban land burned, though our goal in this work is not to improve upon WUI and urban fire modeling tools.

### Fuel Types

All fuel modifications in our simulations were based on the LANDFIRE Fire Behavior Fuel Model 40 (FBFM40), which we selected as our baseline fuel model layer in FARSITE [24]. We tested NB1, TL1, TL2, and a custom urban fuel type in replicating the Tubbs Fire [25]. We used NB3, TL1, and Anderson 2 in buffer testing [26]; to improve accuracy when replicating the characteristics of a banana buffer, we generated a custom banana fuel model using BehavePlus [27].

### Banana and Vineyard/Orchard Fuel Types

We considered two possible edible fire buffers in our primary experiments: banana and vineyards/orchards. Vineyards and orchards are already known to be viable across much of California. We computed spatial suitability maps for banana at present and for future climate scenarios (Figure 1). Bananas are already widely suitable in California, especially along coastal hills that represent the majority of WUI and the region with greatest population, and their suitability will improve as warming proceeds through the century.

**Fig 1.**
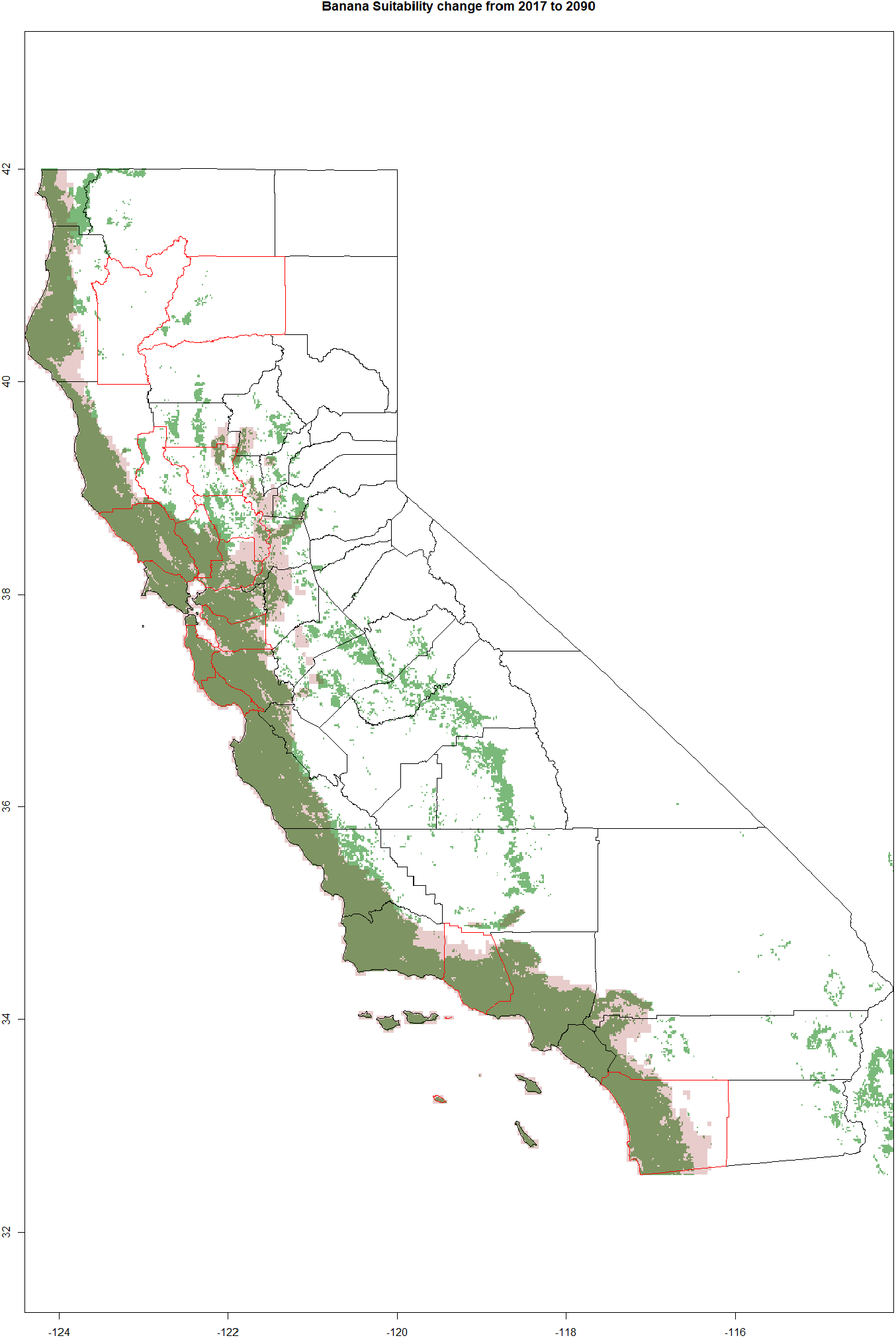
MaxEnt model of banana suitability in current (green) and in 2090 (pink). The model from 2090 represent the mean suitability from eight GCMs (BCC-CSM2-MR, CNRM-CM6-1, CNRM-ESM2-1, CanESM5, IPSL-CM6A-LR, MIROC-ES2L, MIROC6, MRI-ESM2-0) under the RCP 8.5 scenario. Counties outlined in red represent the counties that have had the 10 most intense fires (in terms of damage) in California history. Currently 91 percent of California’s population lives in counties that contain land suitable for banana cultivation.

### Climate Change Scenarios

We consider the effectiveness of fire buffers under two future climate scenarios in 2090 in addition to the 2007 Tubbs Fire baseline. The first is the no mitigation business-as-usual scenario, Representative Concentration Pathway (RCP) 8.5, and the second is the moderate mitigation scenario, RCP 4.5 [28]. We use the average of eight future climate models under each scenario [29] to explore the suitability of banana in California. In modeling future fires, we explore the increase in temperature and the changes in humidity that would occur. Using the mean of eight models provided a way to explore the variation in future predictions to get a better understanding of the efficacy of potential mitigation strategies. We model the temperature and humidity in these future scenarios in the location of the Tubbs Fire.

### Fireline Intensity

Fire buffers can be deployed on the land in a variety of spatial configurations. We consider simple, non-optimized rectangular fire buffer configurations placed on the human-managed wildland that abuts the WUI; optimized spatial configuration may improve the effectiveness of the fire buffers. We define the width of the buffer to be the minimum distance from the wildland side of the buffer to the side of the buffer that abuts the WUI; we configure buffers across a range of sizes. We limit the width of buffers to sizes of agricultural parcels in the region, though even larger buffers have the potential to produce improved fire mitigation. Our canonical buffer for these experiments is a medium-sized 633m buffer (tested alongside a very small 200m buffer, small 390m buffer, and 1280m very large buffer), and we consider a variety of fuel types, including a fully non-burnable buffer, baseline regional vegetation (e.g., a mix of grasses, shrubs, and trees), vineyards/orchards, and bananas.

Fireline intensity with buffers ranging from small (easily adopted) to very large (difficult to adopt) is shown in Figure 2a. We show the intensity of the fire without buffers, with medium-sized vineyard buffers, and with banana buffers from very small to very large, considering the 2017 climate conditions of the Tubbs Fire and projected conditions in 2090 under RCP 4.5 and RCP 8.5 [28]. We also compare against a control and various fuel treatments and their mitigations, such as mechanical thinning and prescribed fire, in a similar environment as studied by Stephens and Moghaddas [30]. We find that a medium-sized banana buffer results in a 96% decrease in fireline intensity at the WUI, comparable to the combined effect of prescribed burns and mechanical thinning.

**Fig 2.**
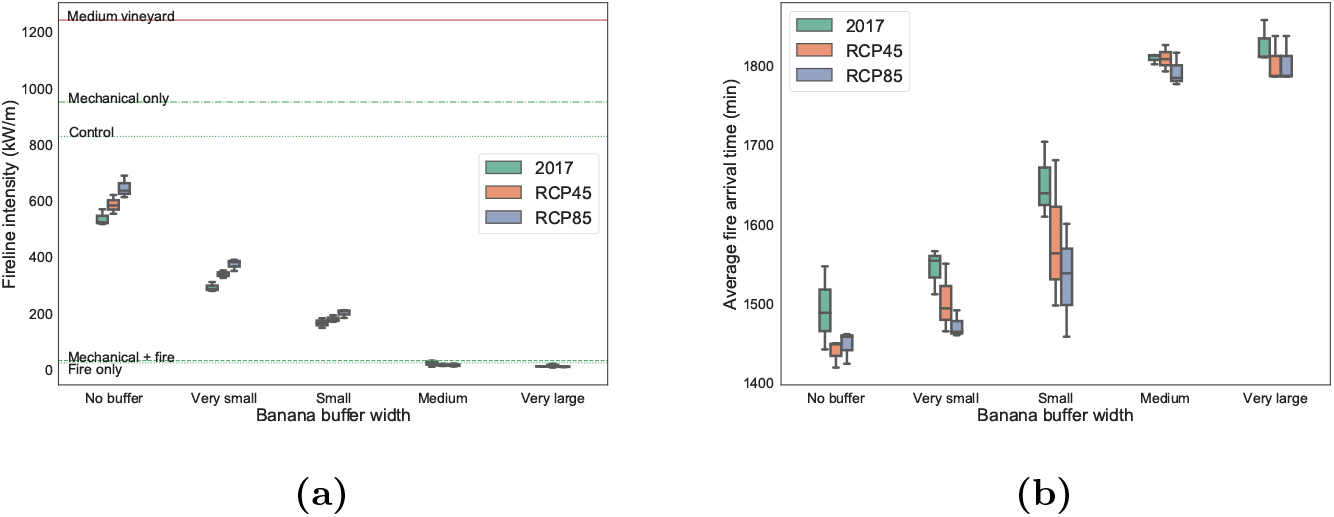
Comparison fire buffers sizes and alternative fire buffers. (a) Comparison of fireline intensity with no banana buffer or with a range of banana buffers ranging from very small to very large, considering the 2017 climate conditions of the Tubbs Fire and projected conditions in 2090 under RCP 4.5 and RCP 8.5. The solid red line shows the fireline intensity of a medium-sized vineyard under 2017 conditions. Dashed lines in green show the intensity given various fuel treatments and their control in a similar environment as studied by Stephens and Moghaddas under 90th percentile weather conditions [30]. (b) Comparison of arrival time of the fire to the side of the human-managed wildland fire buffer region that abuts the WUI, again considering 2017 conditions and 2090 climate projects under RCP 4.5 and RCP 8.5. The y-axis depicts the time since the fire’s ignition. Boxplots show the range of values for five simulations of the stochastic fire model for each buffer.

In Figure 2b we show the time of arrival of the fire to the edge of the wildland edible fire buffer that abuts the WUI. We consider each size of buffer and once again consider 2017 climate conditions of the Tubbs Fire and projected conditions in 2090 under RCP 4.5 and RCP 8.5. We find that very small buffers (≤ 200m) are inadequate to substantially slow the spread of the simulated Tubbs Fire, regardless of buffer fuel type; we performed repeated experiments that indicate that this inadequacy of small buffers is due to ember cast, as winds carry embers over small buffers. The only other buffer type that showed a similarly poor effect is vineyard/orchard buffers at the end of summer, which are are no better and sometimes worse than the baseline fuels and thus do not provide a fire mitigation benefit. We find that medium-sized banana buffers would slow the arrival of the Tubbs Fire by 316 minutes. This would double the amount of time to make a substantive intervention (e.g., initial attack [31]) on a fire. As the time and intensity of the fire are two of the key determinants of the ability of fire crews to stop a fire, this would have major practical impact on the protection of communities. While under climate change the fire spreads faster, we find banana fire buffers continue to have a substantial protective benefit.

### WUI Burn Area

Further, we show that medium-sized and larger banana buffers provide a substantial fire mitigation effect on WUI/urban fire spread. As we mentioned earlier, while FARSITE [21] is not ideal for measuring WUI/urban fire spread, there exists no better alternative simulator for such projections and our validation experiment showed that it is possible to compute an accurate fire perimeter when recreating the Tubbs Fire; we report the values with that caveat.

We find that the medium-sized banana buffer results in an WUI/urban burn rate of 56.9% of the unmitigated baseline. This compares favorably to a vineyard buffer of the same size, which provides no clear protective effect (Figure 3a). Very large buffers (more than a km in width) are even more effective but likely impractical to manage. We also show the findings of fire buffers under future climate change (Figures 3b and 3c). We find that banana fire buffers continue to be effective in these future climate scenarios, in which there is a baseline, no-buffer urban area burn rate by 2090 of 154.8% under RCP 8.5 and 135.0% under RCP 4.5 relative to baseline (2017). We find that a medium-sized banana buffer at this WUI results in a reduced urban burn rate, even under predicted climate change, of 120.1% under RCP 8.5 and of 85.9% under RCP 4.5 relative to the same baseline (2017).

**Fig 3.**
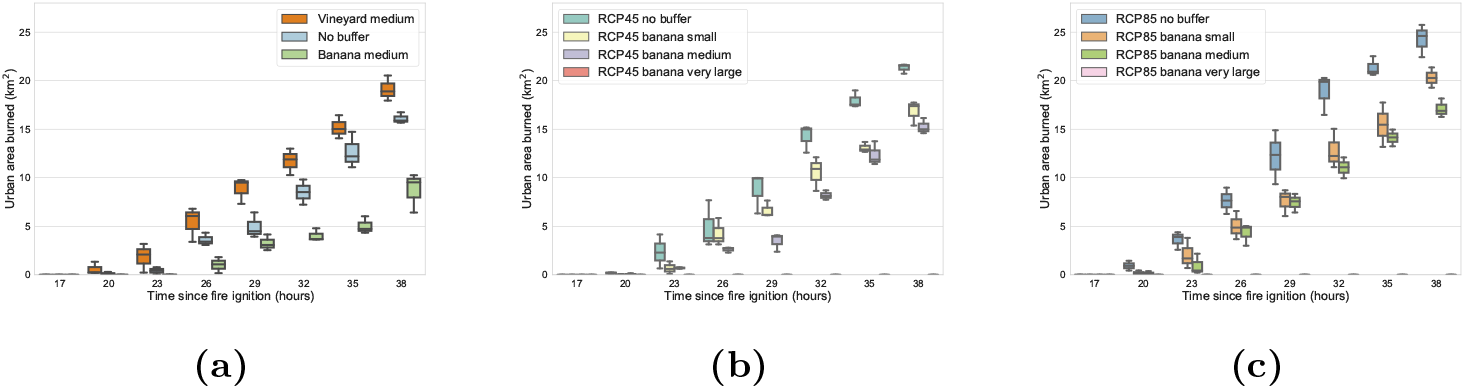
Comparison of fire buffers’ effectiveness in mitigating urban damage in present and future climate scenarios for the Tubbs Fire (2017). Boxplots show the range of values for five simulations of the stochastic FARSITE fire simulation for each buffer. (a) Comparison of urban burn area of baseline (no buffer) scenario and medium-sized vineyard and banana buffers. (b) Comparison of urban burn area in banana buffer of different widths in projected RCP 4.5 2090 conditions. (c) Comparison of urban burn area in banana buffer of different widths in projected RCP 8.5 2090 conditions. In both (b) and (c), very large buffers, which are over 1km in size, entirely stop fire spread.

### Economic and Ecological Analysis for Edible Fire Buffers

Human-managed wildlands have multiple possible uses; for edible fire buffers to be adopted they likely need to create additional revenue while helping to mitigate fire and/or providing additional ecosystem services. To explore this potential value, we studied the cost of planting and harvesting banana by creating an enterprise budget, with costs chosen based on the California context for organic production with recycled water, and yield projections based on banana grown in similar Mediterranean climates. We consider multiple yield, crop value, and land cost scenarios. Based on this budget the potential profit for banana buffers ranges from a low-yield, low-value, high-land-cost scenario with negligible profit (but no loss) to a high-yield, high-value, low-land-cost scenario with profit of 76,136 USD / hectare (see Supplementary Table 1), accounting for planting, irrigation, inputs, maintenance, harvesting, and other relevant costs. This analysis is based on pricing of normal cultivars grown under organic conditions, and does not account for secondary benefits such as a decrease in firefighting or insurance costs or an increase in ecosystem services. There is a potential increase in value if markets develop for speciality cultivars [32]. Even if individual buffers do not achieve the high-yield scenario, any value rather than a liability creates a potential to consider these buffers as a near-term financially- and ecologically-sustainable solution.

In addition to the value of the food produced, there is substantial improvement in land value by changing the groundcover. Aesthetic preferences show that green spaces in and around urban areas, particularly those with trees, are preferred [33]. The desire to have landscapes produce more than one ecosystem service is becoming increasingly important to all aspects of the constructed environment [34]. Municipalities invest in developing public areas that provide multiple ecosystem services; these public areas include parks, roads, schools, and business parks. Public firebreaks that also provide healthy, nutritious food could provide profit, savings in terms of fire damage, and additional jobs, creating win-win scenarios.

## Discussion

### Building and Land Use

The last two decades have seen legislative efforts to mandate and to reimburse individual homeowners and municipalities to respond to fire risk [35]. The increase in fires has led to changes in building materials [36] and new regulations for WUI areas. These regulations have helped change patterns of vegetation management through increased clearing of dead plant material and by limiting the use of highly flammable plants [37]. We discuss current California land use in supplementary materials and note the current human management of wildlands that abut the WUI.

### Fuel Reduction

Fuel reduction is a key practice in fire risk management. The U.S. Forest Service performs spatial risk analysis by examining simulated fires given a range of hypothetical fuel treatment scenarios, including change of the canopy cover and fuel map. However, the actual costs and complexity of such treatment makes it difficult for land managers to reduce both current and future fire risk [38]. Similarly, for forest fuel-reduction treatments such as prescribed fire and its mechanical surrogates, the effect is transient so long-term mitigation is hard to achieve [39].

### Firefighting Resources

The predicted increase in fire occurrence and severity will decrease the success rate of initial attack making fire control more difficult and necessitating an increase in firefighting resources [40]. The 3–5 hours of initial attack is considered to be crucial for mitigating fire spread and suppression costs [31]. Suppression efforts at this stage may include air tankers and control line resources such as pumps, hoses, bulldozers, and shovels [41]. Initial attack fails when suppression resources arrive too late, fire intensity is too high, or when firefighters fail to contain spread to a predetermined target area [42]. If initial attack is insufficient and an advancing fire crosses a defined evacuation trigger, then an evacuation order is released to nearby communities [43]. Reducing the rate of spread of fires during initial attack could reduce the need for evacuations, mitigate evacuation risk, and allow firefighting personnel to execute substantive suppression efforts.

### Firebreaks and Fire Buffers

Creating firebreaks through prescribed burns or mechanical methods has also proven to be effective. Recent work has proposed multiple strategies, including: 1) replacement of the species mix to less flammable natives [44], 2) ecological vegetation management in high-risk areas such as by utility companies on lands near power lines [45], 3) conversion to low-growing succulents, irrigated agriculture [15], or other green firebreaks [46]. In addition to pre-constructed firebreaks, firefighting techniques incorporate the creation of bulldozer lines during initial attack [47]. However, each of these strategies imposes up-front and ongoing costs.

### Practical Considerations for Edible Fire Buffers

While we have focused on banana as a key crop for edible fire buffers, next we discuss some requirements for and limitations to its use.

#### Microclimatic suitability

Not all Mediterranean areas are appropriate for banana cultivation, though present WUI-adjacent areas in California are nearly all suitable for banana cultivation as they frequently occur in thermal belts with little to no frost, so with careful cultivar selection, bananas are likely widely appropriate. Additionally, due to the Mediterranean climate it appears that buffers need to be larger than in regions such as China where green firebreaks are employed [46]. In supplementary text we also consider alternative crop species for edible fire buffers.

Banana is a high-water-need crop; we find through an informal analysis in California that recycled water widely available to meet this need. In addition, many high-fire-risk WUI areas have been developed more recently, and these areas have ubiquitous recycled water, currently used for ornamental landscaping. In addition, banana buffers could leverage even lower-cost, undisinfected secondary recycled water [48]. Recycled water also carries with it plant nutrients that can decrease or eliminate the need for supplemental fertilizer. As California is a region with increasing hydrological volatility and long dry seasons, there is a need to put available water resources to better use. Maximizing reuse of municipal water, in the form of recycled water, to grow food while also protecting those same WUI areas from fire is a virtuous cycle.

#### Groundcover and establishment

The high-fire-risk areas we consider are often hilly and covered in fire-prone vegetation. While banana plants themselves are fire resistant, the land underneath the banana canopy must be managed as well. Annual dryland-adapted grasses quickly cover such WUI lands, sometimes within a single season after a fire.

Such grasses would significantly decrease the effectiveness of any buffer or break. Thus, edible fire buffers should be established along with low-growing, non-flammable groundcover species; common examples include *Senecio mandraliscae* and low-growing cultivars of *Aloe ciliaris*. Such groundcovers can be established along with the crop species with a cellulose-based biodegradable weed fabric to prevent emergence of grasses during establishment. To avoid a monoculture of banana buffers that succumb to the spread of a pest or pathogen (e.g., Panama Wilt), thereby undermining the fire buffer, deployments of banana buffers should employ spaciotemporal diversity of cultivars. Such diversity is also likely to be beneficial to adapt to local microclimatic needs and market demand. Testing multiple groundcovers is feasible because unlike most horticultural crops, upkeep on banana farms is minimal [49].

#### Land considerations

Soil suitability for banana cultivation is a possible concern, but in the regions we consider, soils are often well-drained clay loam (e.g., the regions of concern in California, including the area of the Tubbs Fire), with good suitability for banana cultivation with water and fertility supplementation (e.g., through recycled water). If the land has recently suffered a burn there is a risk of soil hydrophobia that exacerbates runoff; such soils would benefit from limited-depth mechanical tillage before banana cultivation begins. Pathogenic nematodes, fungi, and viruses for banana are currently not present in California and thus not a factor in this context, but may need consideration in other regions. The major limitations to banana cultivation are not biological; banana grows well on hillsides and in erosion-prone soils and banana root systems are well adapted to shallow soils [49], yielding potential erosion-control benefits. Rather, zoning (e.g., residential, agricultural, parkland) and political authority (e.g., cities, counties, HOAs, utilities) are likely to present challenges. The primary purpose of edible fire buffers for fire mitigation is in line with existing land uses and zoning in human-managed wildland areas that abut the WUI. The availability of recycled water on a large scale has transformed the feasibility of novel land use change proposals such as edible fire buffers.

#### Extreme Fire Behavior

Recent years have seen a rise in “extreme” fire behavior, including more frequent occurrence [8] of lightning-producing pyrocumulonimbuses and pyrovortices [50]. It seems unlikely that any buffers or breaks – even if completely non-flammable – can stop the spread of such fires. However, it may still be the case that fire buffers, in combination with other mitigation efforts such as low-flammability urban landscaping and building materials, can meaningfully respond to worsening extremes. Further, katabatic “Santa Ana”-type winds have led to increased severity in many of the most damaging effects. Practical, sustainable options that improve the use of water, provide food, improve aesthetics, and protect people and homes are needed for municipalities to take concrete action to change the status quo; edible fire buffers are one such response. Further, the increasing fire burden under future climates [2] necessitates creative solutions and edible fire buffers provide the potential for substantial relief under both RCP 8.5 and RCP 4.5, showing how creative combinations of climate science, ecology, and agriculture can tackle wicked problems. Looking at the historic use of banana in California, its future suitability, and the value such edible fire buffers would yield, banana becomes a prime test case for how to use edible fire buffers to sustainably mitigate wildfire risk.

### Alternative Buffer Scenarios

We explored banana in depth as a possible crop for edible fire buffers, but it is not the only possible option. Other low-flammability, high-value crops could appropriately mitigate fires and serve as fire buffers. The key properties of such fire buffers are that they are not only low-flammability and yield a net profit, but also that they are suitable in other dimensions for the zoning, terrain, and context of many WUI lands. Specifically, standard vegetable crops (e.g., of leafy greens or vegetables), while low-flammability, are not suitable given the need for significant labor and machinery, and the need for relatively flat terrain. Many WUIs, especially in California, have variable terrain and are in an environment (e.g., residential neighborhoods) where large machinery and the workforce of a working farm are not suitable. Banana, like most perennial crops, requires less labor and is more adaptable to varied terrain. In addition, automated systems could be developed to automate aspects of the banana harvest; such an environment would provide a prime location to test such new technology. Thus perennial, low-flammability species are ideal.

Ginger is a low-flammability culinary crop that would be suitable for growing in the same regions as banana, though mechanical harvesting may be required to keep harvest costs low. Indeed, there is a long history of agroforestry in banana orchards, and many different cover crops have been successful [51]. Including an intercrop such as ginger could increase the value of the total crop, take advantage of microclimates (as ginger grows well in partial shade and is low growing), and increase species diversity in the buffer region.

The most widespread high-value crop in Mediterranean climates is wine grapes. We evaluated the effectiveness of vineyards as fire buffers, and found that they may not be suitable due to their flammability. While currently not a widely grown crop, carob is a potential high-value, high-yield, and low-flammability crop appropriate for high-fire-risk Mediterranean climates. There is some evidence that carob is less flammable than other similar dryland species [20]. Since carob can grow without supplemental irrigation in dry-summer regions, it may be appropriate in settings where irrigation is either unavailable or too costly.

Finally, some regions may consider physical non-flammable fire buffers, such as ones made from concrete or metal, as opposed to crop-based buffers. However such buffers would impose a high installation cost, would require annual maintenance (to keep soils from building up on top thereby enabling grass growth), would provide no revenue or ecosystem services, and would exacerbate problems such as water and soil runoff. Like bulldozer lines, they are an expensive and intrusive intervention that may be suitable in certain circumstances but unlikely to yield win-win outcomes.

### Current California Land Use

It is also important to consider pre-existing land use where edible fire buffers are considered. Unlike truly wild areas, the lands we consider for such buffers in California are human-managed wildland regions that are not truly wild due to current fire management practices, development, and proliferation of introduced annual grass species that have outcompeted native perennial grasses subsequent to severe overgrazing [52, 53]. Banana fire buffers can provide ecosystem services similar to existing human-managed wildland areas that abut the WUI [51, 54]. In addition, future climate conditions with increased fire frequency and severity are projected to induce losses in ecosystem services; these can be mitigated by reducing fires through reduced greenhouse gas emissions [55] and possibly, in human-managed wildland areas, through the creation of edible fire buffers.

### Development of fire-resistant cultivars

Plant moisture content is a trait that appears to have genetic variance within species and can be selected for [56]. Thus breeding programs can identify species and populations within species that that could be used for edible fire buffers. This could be a fruitful use of public funds: to select new, multi-functional varieties.

## Methods

### Overview

In Supplementary Figure 1 we depict the workflow for our experiments and also for a potential deployment of edible fire buffers on the land. Our experiments use canonical tools for fire behavior modeling. We use FARSITE [21] for fire simulation, LANDFIRE [24] for landscape data, and BehavePlus [27] for fuel type modification. We use standard GIS tools such as ArcGIS [57] for buffer placement and GDAL [58] and rasterio [59] for rasters. We use seaborn [60] and geopandas [61] for data processing and visualization. We use GeoMAC [62], NOAA [63], and Silvis Lab WUI data [64].

### Replicating the Tubbs Fire

We acquired a Tubbs Fire base map from LANDFIRE, which provides country-level fuel maps for use in FARSITE and FlamMap [24]. Additional landscape data was obtained from the LANDFIRE Data Access Tool (LFDAT) [65]. To accurately represent weather conditions, we supplemented our existing NOAA weather data with wind speed data from news reports, as extreme weather was captured with higher fidelity by local management agencies than by NOAA stations (which suffered from power outages) [66]. We performed preliminary calibrations on our parameters based on standard guidelines for fire behavior modelling [21]. FARSITE incorporates existing models for surface fire [67], crown fire [68], point-source fire acceleration, spotting, and fuel moisture [21, 69]. Additionally, FARSITE uses weather and wind inputs, so we were able to incorporate future weather scenarios into our simulation.

#### Determining the Optimal Urban Fuel Type

To define the WUI in our study area, we followed the description set forth by SILVIS Lab’s WUI dataset, as well as a standard risk assessment framework that characterizes WUIs as HVRAs (highly valued resources and assets) [70]. We classified urban areas in Santa Rosa based on housing density of above 20 housing units per square kilometer, and negated all vegetation that contributed to fire risk [71]. Urban boundaries were based on the US Census Bureau block data, following our definition of WUI [6]. CalFire 2013–2017 housing damage data [72] was used to assess infrastructure damage and validate our definition of protected area. Our classification of urban areas was applied towards determining an urban fuel type which most closely modelled the satellite data obtained from USGS. We compared the urban fuel type from the LANDFIRE base map, NB1, against several other standard fuel models [25] and a custom model, and calculated the F1 score of each to assess the accuracy of our result against the ground truth. TL2 was determined to best represent fire propagation in urban areas.

### Banana Fire Buffer Modeling

In order to identify the suitable area for banana cultivation in California, global banana occurrence points were downloaded from Global Biodiversity Information Facility [73]. These occurrence points were used as input for MaxEnt [74] suitability modeling, which was implemented from geo-referenced coordinates implemented in the software R, under current climate conditions using 19 bioclimatic variables [29] and under future conditions representing RCP 4.5 and RCP 8.5 [28]. Future models were constructed using eight GCMs (BCC-CSM2-MR, CNRM-CM6-1, CNRM-ESM2-1, CanESM5, IPSL-CM6A-LR, MIROC-ES2L, MIROC6, MRI-ESM2-0) at a 2.5 arc minute resolution in 2090 [29]. Suitability maps of current and future models were overlaid to explore which counties have potential for cropping interventions. Suitability models were considered accurate if they complied with the following conditions: (i) five-fold average area under the test ROC curve (ATAUC) is greater than 0.7, (ii) the standard deviation of ATAUC (STAUC) is less than 0.15, and (iii) at least 10 percent of grids for each model has standard deviation less than 0.15 (ASD15).

#### Preliminary Buffer Testing

To test the potential of food crop firebreaks for minimizing fire damage, we constructed buffers with fuel type NB3, which is classified as non-burnable agricultural land [25]. Keeping all other baseline parameters constant in FARSITE, we simulated fire behavior with buffers of the following widths placed on human-managed wildlands that abut the WUI: 200m (very small), 390m (small), 633m (medium), 1070m (large), and 1280m (very large); for clarity of the figures we omit large buffers. Width values were determined by creating buffers in order of increasing size, with exact value selection constrained by the granularity of the raster in ArcMap; this yielded values that are distributed across the range of agricultural land sizes that are standard in the region.

#### Validating the Custom Banana and Vineyard/Orchard Fuel Models

Of the non-spatial fire behavior modeling systems available, we selected BehavePlus [27] to evaluate the behavior of our custom fuels. We began with TL1 [25] as a baseline fuel model (as it is characterized by low spread rate and low flame length), and then we modified fuel load, fuel bed depth, and fuel moisture parameters to model banana crops. Specifically, bananas have a range of total moisture from 76–92% depending on drought conditions [19], which, converted to a dry weight basis is a minimum 316%, higher than the upper cutoff allowed by BehavePlus (which was designed for wildland species that rarely have such high moisture content); thus, we use the maximum value for live fuel moisture. Similarly, for dead fuels, the banana buffer is highly managed so little to no dead fuels are to remain and the groundcover will be of a succulent plant species. However, should management fail to note dead banana plants, we can account for the dead fuels based upon studies of banana plant drying, which finds once again that bananas will yield dead fuels of higher moisture than BehavePlus’s maximum settings [75], as banana is a non-woody plant species with very high water content. We compared simulated fire behavior between our banana fuel type and agricultural NB3, and found that the two yielded similar results. We tested buffers of varying widths under NB3 and our custom banana fuel model. Bananas behave similarly to NB3, and for both fuel types, fire spread slows as buffer size increases.

To validate our results, we performed similar experiments using a conventional vineyard/orchard buffer, which we hypothesized to be less fire-resistant than bananas. As well-studied cropping systems with proven economical benefits, vineyards and orchards provide a comparable model for banana buffers as a profitable land use for a potential fire buffer. Based on previous work assessing land covers, the best match to the vineyard and orchard landscapes is Anderson 2, which was used to describe tree crops – including vineyards and orchards – in Sardinia, Italy [76], which has a similar climate to California. Both banana buffers and vineyards/orchards were modeled with medium buffer widths as this width demonstrates moderate fire mitigation effects. In addition to this validation in simulation, we note that vineyards in recent California fires have performed poorly as fire breaks, often burning substantially; this is unsurprising as vineyards seldom have supplemental irrigation applied late in the season and are thus dry and woody. Banana orchards are not common today in California, but there are reports an Australian banana orchard with grass groundcover burned during a major recent wildfire, damaging the banana leaves but leaving pseudostems intact to resume growth soon after the fire was extinguished; this points to the importance of our incorporation of non-flammable groundcover.

When reporting fireline intensity and arrival time for banana buffers, we report values directly from FARSITE as produced in our simulation runs. We compute the mean of the reported fireline intensity for the pixels of the buffer region in each run of the simulation. Similarly, we compute the mean arrival time of the fire, in minutes from ignition, across the pixels of the buffer region in question in each simulation run.

#### Validating Ember Spotting Behavior

To ensure that the fire simulation accurately reflects the extreme ember spotting behavior seen in recent WUI fires in California, we ran several validation experiments with and without fire buffers and with and without ember spotting enabled in the simulator. With ember spotting enabled, lowering the wind speed below the actual historical wind speeds also showed lower fire spread, as expected.

#### Climate Scenarios

We used RCP 4.5 and RCP 8.5 to simulate future scenarios [28]. We acquired temperature projections for the study area from eight GCMs (BCC-CSM2-MR, CNRM-CM6-1, CNRM-ESM2-1, CanESM5, IPSL-CM6A-LR, MIROC-ES2L, MIROC6, MRI-ESM2-0 [29]) and projected the temperature for Tubbs Fire scenarios. We calculated the mean temperature offset based on the maximum difference between each predicted temperature and the 2017 Tubbs Fire area data from October 2017. We considered ocean warming as a contributing factor to decreases in relative humidity on land [77]. After projecting the temperature, we modified the local relative humidity value for simulation of fire spread based upon the static dew point, yielding, in future scenarios, lower humidity that drives faster fire spread. Fire progression under the no buffer scenarios for projected climatic conditions shows that the urban burn will increase. We compared fire spread under RCP 4.5 and RCP 8.5 [28].

### Economic return on edible fire buffers

Fire buffers of any type are not widely deployed today and are not currently financially sustainable. We computed the cost of edible fire buffers, independent of their fire mitigation savings, using an enterprise budget modified from the University of Florida banana template [78]; the modifications are shown with references for changes that were made in order to develop a range of potential buffer value scenarios in Supplementary Table 1. We consider a range of yield scenarios, banana fruit value scenarios, and land cost scenarios. All cost values are given in 2021 USD.

#### Practical considerations for conversion of land in California WUI areas

Converting existing WUI land to edible fire buffers is likely to be an intricate and site-specific process that cannot be fully addressed here. In brief, our consideration was whether such conversion is practically feasible, but we leave to future work the exact procedures by which such conversion can or should be done. In this, we take a perspective gained from our professional experiences in agricultural and horticultural research and practice in Mediterranean and tropical regions and successes in banana cultivation in both Northern and Southern California. WUI lands suitable for edible fire buffers are largely covered by low- to medium-density annual grasses and low-growing shrubs. Some number of these regions are beginning to type convert as decades of fire suppression and climate change lead to a greater frequency and intensity of fires. In addition, these lands are often in an already-disturbed state given the very urban lands / housing that we aim to protect. Establishment of edible fire buffers on these lands is likely largely limited by infrastructure availability, particularly water for irrigation.

## Addendum

### Data availability

The data that support the findings of this study are available in the databases listed in the Methods. Data and code are available from the authors upon request.

---

**Supplementary Figure 1.**
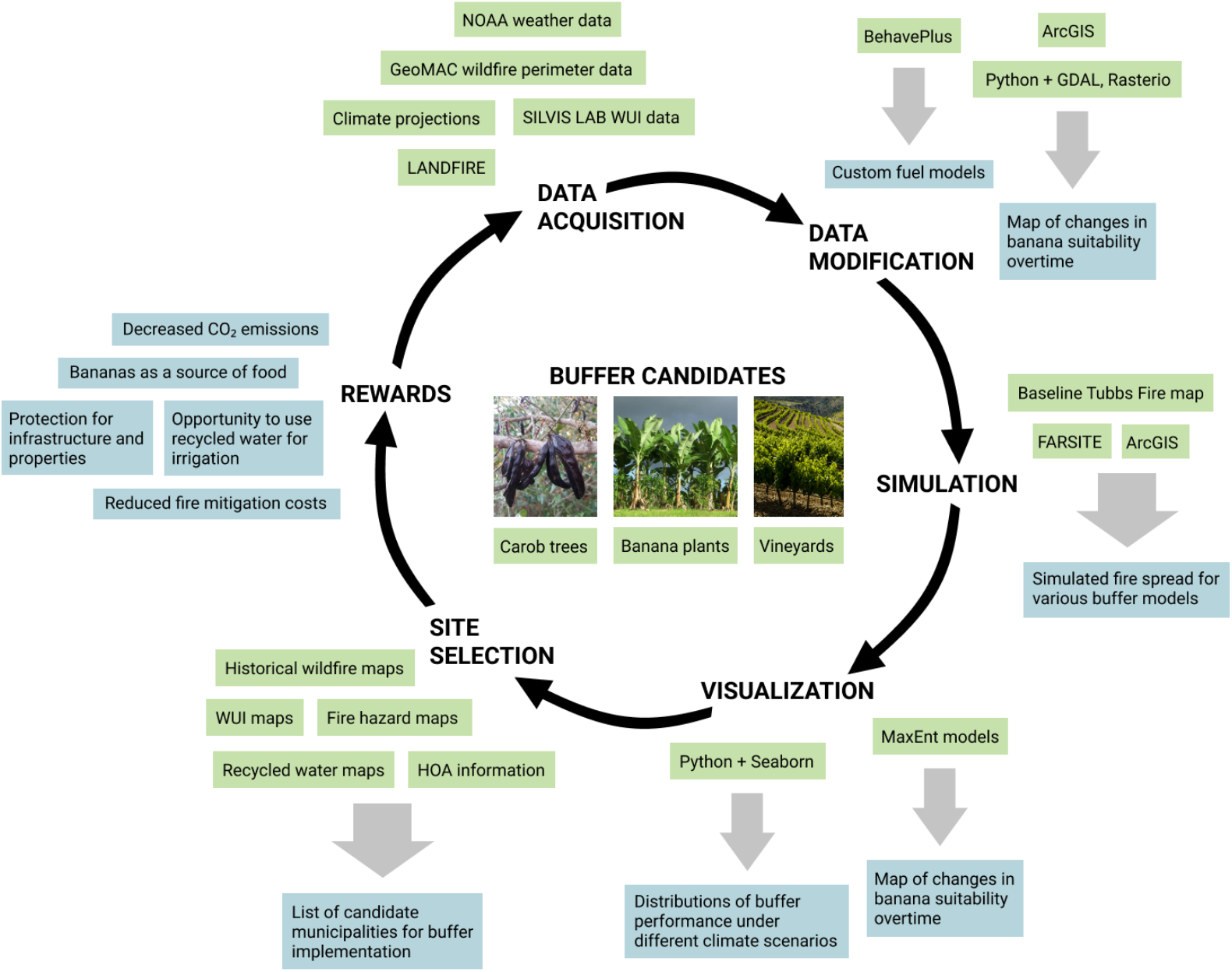
Workflow for modeling and analysis of Edible Fire Buffers using existing software tools and datasets, including post-simulation considerations such as site selection and benefits.

**Supplementary Table 1.**
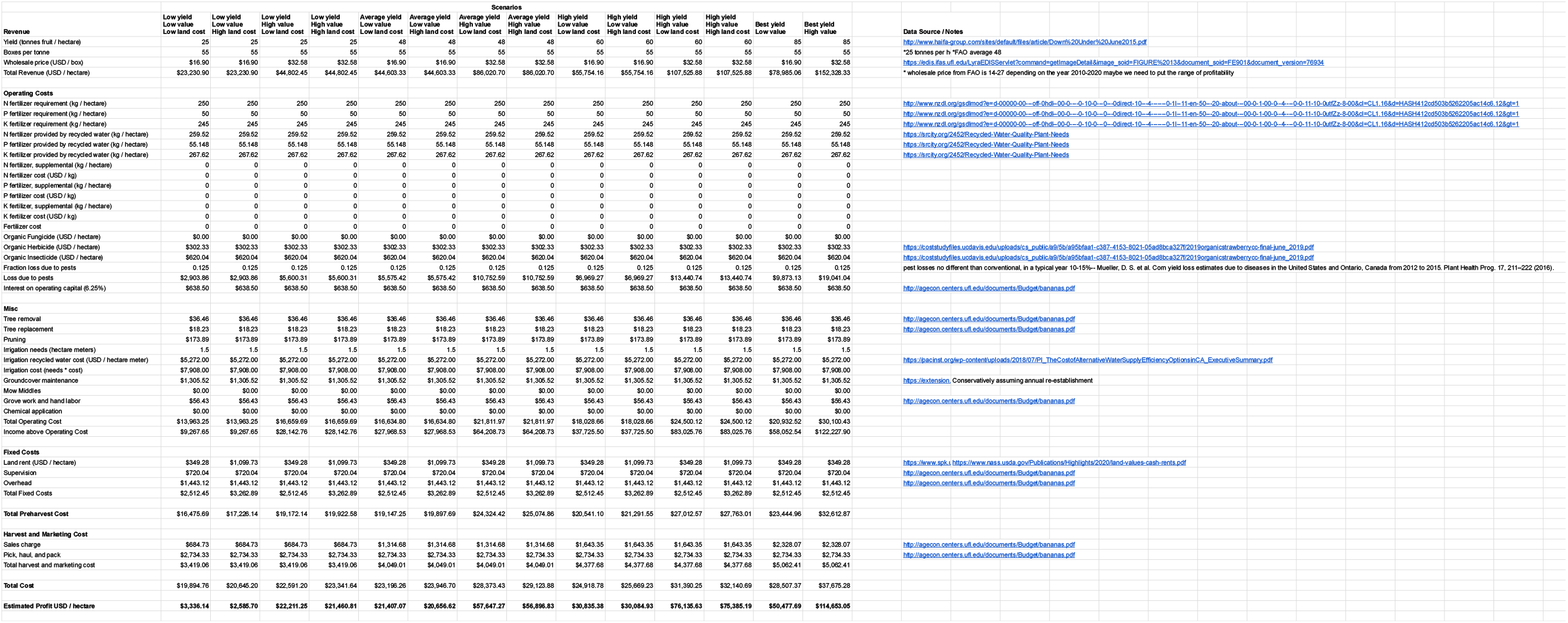
Enterprise budget for banana fire buffers in California considering low, average, high, and best case yield, low and high crop value, and low and high land cost scenarios. All values are given on an annual basis. All costs and revenues given in 2021 US Dollars.

